# Learning protein function through autonomous experimental interaction

**DOI:** 10.64898/2026.08.14.744985

**Authors:** Coban Brooks, Pascal Notin, Philip A. Romero

## Abstract

Biological AI learns primarily from existing observations, but many questions cannot be answered from available data alone. Here we show that AI can instead acquire knowledge by acting directly on biological systems and learning from the consequences. We developed a closed-loop framework in which autonomous agents design protein variants, construct and characterize them in a robotic laboratory, learn from the resulting experimental feedback, and decide what experiments to perform next. We then allowed the system to operate continuously and without human intervention for approximately one month, during which multiple agents independently explored protein sequence space while learning from shared experimental experience. Applied to glycoside hydrolases, the agents discovered enzymes with substantially altered substrate specificity toward non-native sugars and progressively learned the structure of the underlying sequence–function landscape. The resulting experimental experience also revealed determinants of substrate specificity and protein expression that were not specified as learning objectives. These results demonstrate that AI can autonomously interact with biology over extended periods to acquire knowledge through experience, establishing a framework for biological discovery driven by continuous experimental interaction.

## Introduction

Recent advances in artificial intelligence have transformed biology. AI systems can now model the relationships between protein sequence, structure, and function with unprecedented accuracy, resolution, and scale. Models such as AlphaFold^1,2^, RFdiffusion^3–5^, and ESM^6^ can predict protein structures, infer function from sequence, generate de novo proteins, design biomolecules with tailored properties, and, more generally, provide flexible model scaffolds that can be adapted and fine-tuned for specific applications^7–9^. Their success has been built on more than half a century of systematic data generation across genomics, metagenomics, structural biology, and experimental biochemistry — a collective effort that has produced billions of protein sequences and hundreds of thousands of protein structures^10,11^.

Despite these advances, the current AI paradigm in biology remains largely observational. Most models learn by extracting statistical patterns from static biological data derived from naturally evolved sequences^12^. While highly effective within these observed regimes, they often struggle to engineer proteins with novel or non-natural functions^13,14^. This limitation arises because desired functions are often poorly represented or absent from available datasets and frequently lack sufficient mechanistic understanding to guide rational design^15–17^. More fundamentally, observational learning is constrained to information already present in the data. It can identify correlations and interpolate between known examples, but it cannot actively perturb biological systems to generate new information about how sequence changes alter function. To move beyond these limits, AI must be able to learn through action and experience rather than observation alone^18–23^.

Here we present a closed-loop learning system that acquires knowledge through experimental interaction, iteratively perturbing biological systems, measuring their response, and learning from the resulting feedback. The system couples a generative protein language model for sequence design with a fully automated robotic laboratory capable of gene assembly, protein expression, and biochemical characterization in a continuous experimental loop. At each iteration, the model proposes protein variants, the laboratory builds and tests them, and the resulting measurements are fed back into the model to guide subsequent rounds of design. We apply this framework to glycoside hydrolase enzymes to study substrate specificity and engineer activity toward non-native sugars. As a demonstration, we implement the system as a coordinated multi-agent platform in which three agents pursue complementary engineering objectives toward glucose, xylose, and mannose while contributing observations to a shared experimental memory and common learning model. Across fully autonomous multi-week campaigns, the system discovers novel glycoside hydrolase variants with threefold shifts in substrate specificity toward non-native sugars. More broadly, this work establishes a framework for AI systems that acquire knowledge of biological function through autonomous experimental interaction.

## Results

### An autonomous agent for iterative exploration of protein sequence space

At the core of autonomous experimental interaction is an agent that can act in a laboratory environment (Fig. 1a). The agent proposes experiments, the laboratory executes them and returns measurements, and those measurements are used to update the agent’s internal model and guide subsequent decisions^24–31^. Enzyme function remains among the most important and challenging properties to predict and engineer in biology, making enzyme engineering a natural setting for learning through experimental feedback. The central challenge, then, is deciding what sequence to test next given the agent’s current understanding of the sequence-to-function mapping^32,33,34^. Because protein sequence space is vast and only a tiny fraction can be experimentally evaluated, each experiment must balance exploitation of promising variants with exploration of uncertain regions of sequence space, simultaneously searching for improved function while generating information about the surrounding fitness landscape^35^.

**Figure 1:**
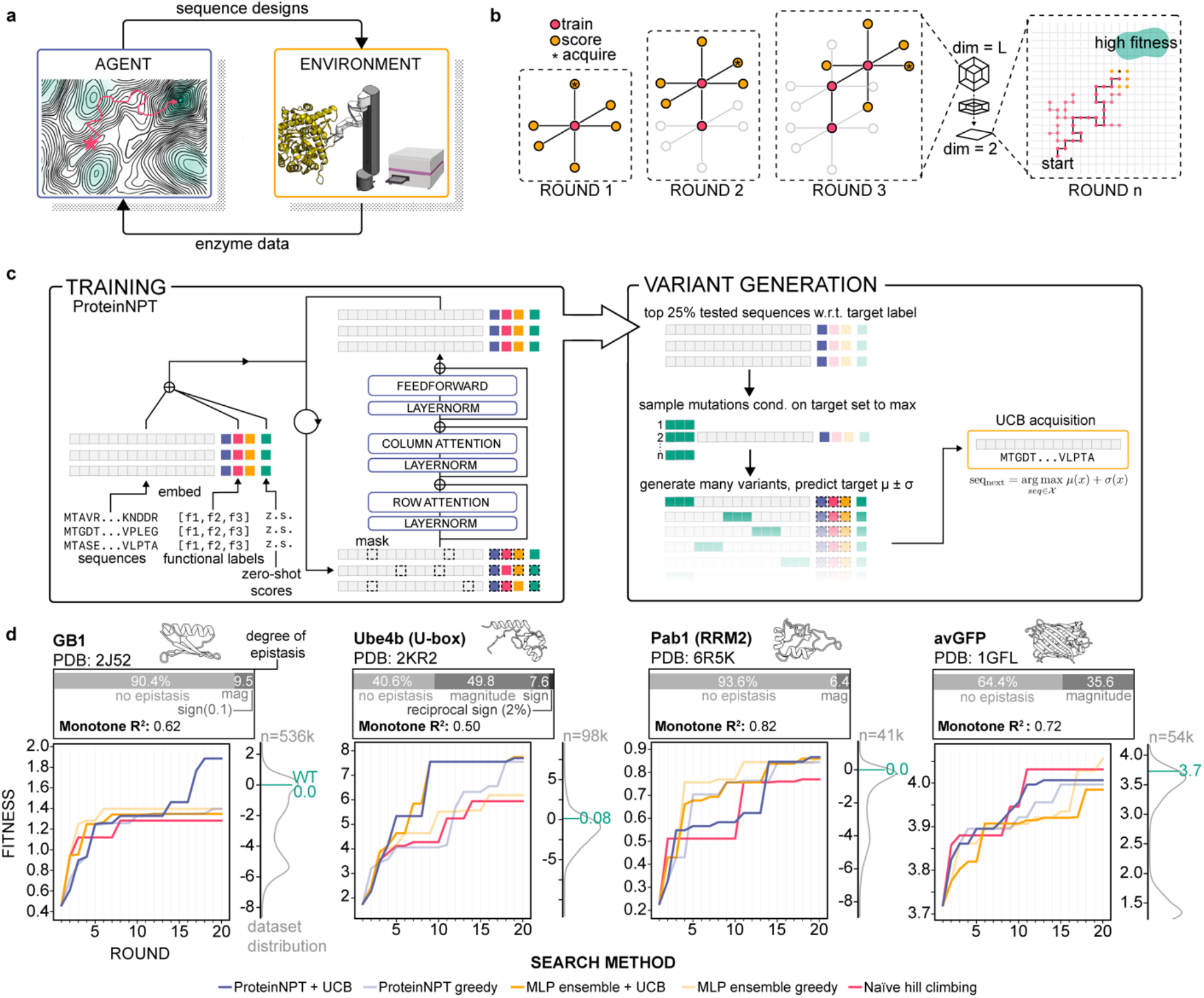
A generative agent for exploration of protein sequence space. (a) Autonomous experimental interaction consists of an agent that selects experiments and an environment that executes them and returns experimental feedback. (b) Protein engineering is represented as an iterative search through sequence space, in which the agent generates and evaluates variants near experimentally characterized sequences while exploring multiple paths toward improved function. (c) Overview of model training and variant generation. ProteinNPT learns sequence–function relationships from accumulated experimental measurements, generates candidate variants by conditional sampling, and selects experiments using upper confidence bound (UCB) acquisition to balance predicted function and uncertainty. (d) In silico comparison of the agent (ProteinNPT + UCB) with four alternative search strategies across four diverse deep mutational scanning datasets, with each dataset serving as the experimental oracle. The top row summarizes the epistatic structure of the high-fitness regions explored during search.

To address this, we developed an autonomous agent that iteratively explores protein sequence space using a generative protein language model (Fig. 1b,c). The model is based on ProteinNPT^36,37^ and learns relationships between sequence variation and function from experimental data while leveraging broader evolutionary priors from pretrained sequence models. At the end of each experimental round, the agent updates its internal model using all accumulated sequence–function measurements, progressively refining its understanding of the local fitness landscape as new experimental feedback is acquired^38^. The agent then uses this learned representation to propose new enzyme variants for experimental testing. High-performing sequences from previous rounds serve as starting points for diversification, from which the model generates candidate variants and evaluates them based on predicted function and uncertainty. The agent then selects experiments using an upper confidence bound (UCB)^39,40^ acquisition function that balances predicted function with predictive uncertainty. Top-ranked candidates are selected and sent to the laboratory for experimental evaluation, creating an iterative cycle in which each round of experimentation both searches for improved variants and expands the agent’s understanding of the protein fitness landscape.

We validated the agent’s ability to select informative experiments using existing deep mutational scanning datasets as *in silico* experimental environments. In this setting, experimentally measured sequence–function data serve as a ground-truth oracle, allowing us to simulate iterative rounds of experimental design while benchmarking search performance under controlled conditions. We challenged the agent to optimize protein fitness across four diverse datasets—Pab1^41^, GB1^42^, Ube4b^43^, and avGFP^44^—spanning diverse structures and functions, fitness distributions, and degrees of landscape epistasis. We compared the agent against four alternative search strategies: ProteinNPT with greedy acquisition that ignores predictive uncertainty and selects only the highest-scoring sequences, an ensemble of multilayer perceptrons with both UCB and greedy acquisition, and random sampling. Each search was initialized with 10 random sequences and continued over 20 iterative rounds of 10 sequence acquisitions each. Across datasets, the agent rapidly identified high-fitness variants with high sample efficiency, consistently reaching top regions of the landscape while only experimentally evaluating less than 0.04-0.49% of the data set (Fig. 1d, Fig. S1).

Although the ProteinNPT + UCB agent performed well across all datasets, its advantage over simpler search strategies depended on the structure of the underlying fitness landscape. The advantage was most pronounced on GB1, where the agent climbed 35–47% higher than the fitness plateaus reached by all other methods. Reaching these high-fitness variants required both ProteinNPT’s ability to model specific epistatic interactions and uncertainty-guided exploration; removing either component caused the search to plateau at substantially lower fitness. On Ube4b, either capability alone was sufficient to improve search: the activating M53L mutation provided an accessible path toward the fitness peak, allowing either ProteinNPT-guided prediction or uncertainty-guided exploration to identify progressively improved variants. In contrast, on Pab1 and avGFP, simpler search strategies performed comparably. Together, these results show that the agent can efficiently navigate diverse protein fitness landscapes, with expressive sequence modeling and uncertainty-guided exploration providing the greatest advantage when reaching high-fitness regions requires navigating epistatic interactions.

### An automated laboratory for generating experimental feedback

To enable autonomous experimental interaction, an agent must be able to perform experiments and observe the consequences of its actions^45^. In protein engineering, this requires constructing designed sequences, measuring their function, and returning experimental feedback rapidly enough to guide subsequent decisions. We therefore developed a fully autonomous robotic laboratory that serves as the experimental environment for the agent. The laboratory consists of a centralized robotic arm connected to a liquid handler, refrigerator, thermal cycler, plate reader, and plate stack, enabling automated movement of materials between instruments and execution of the complete protein engineering workflow (Fig. 2a). Together, the computational agent and automated laboratory constitute PRAXIS (Protein Autonomous eXperimental Interaction System), a closed-loop platform for autonomous experimental interaction.

**Figure 2:**
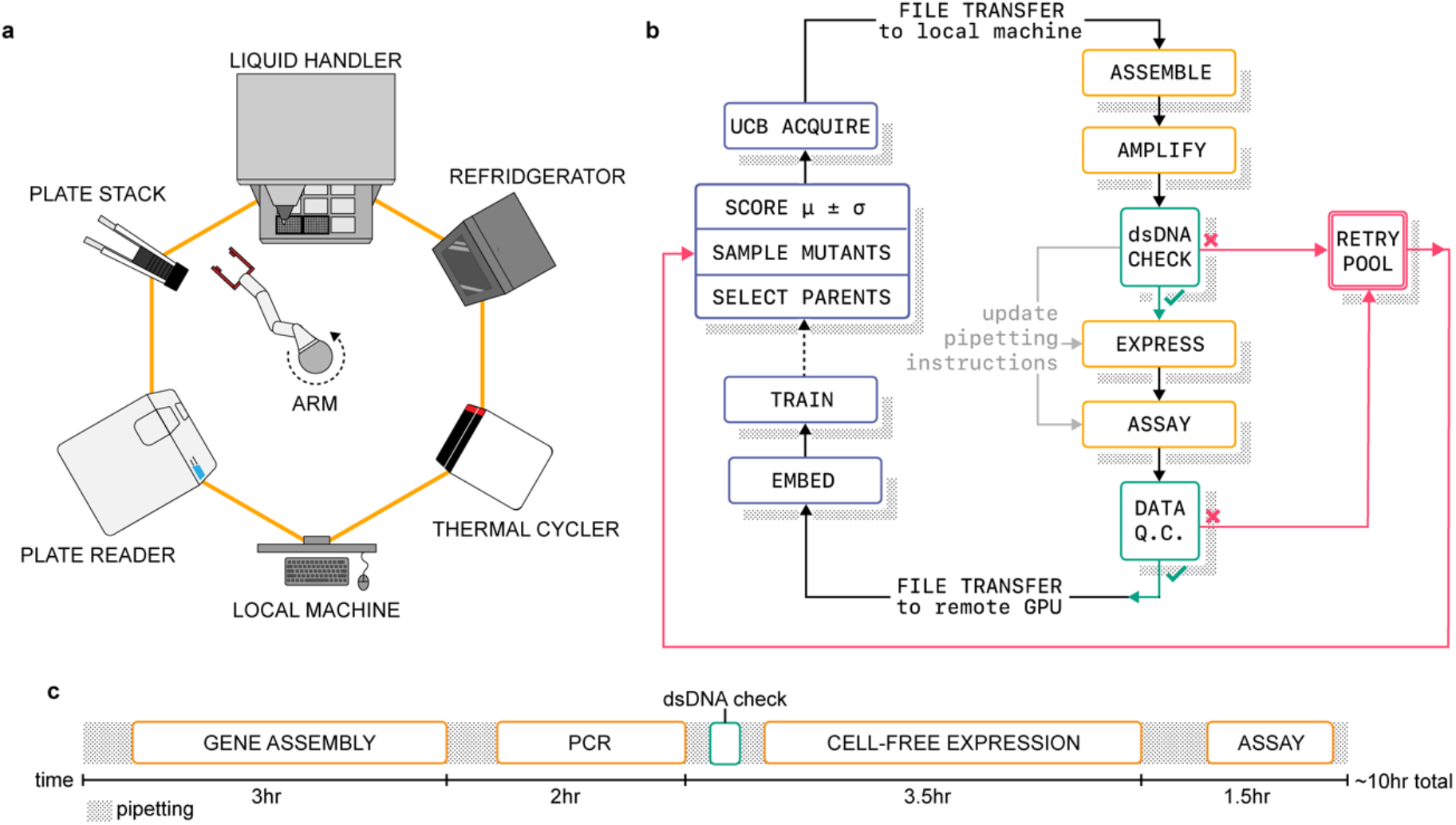
An automated laboratory for generating experimental feedback. (a) The experimental environment consists of peripheral laboratory instruments arranged around a central robotic arm that transfers materials and labware between them. (b) Integration of the computational agent and experimental environment, with sequence designs sent to the laboratory for testing and resulting measurements returned to the agent for model updating. (c) Timeline of a single experimental round, showing the individual reactions and corresponding automated procedures required to progress from sequence design to functional measurement.

To generate experimental feedback, the laboratory receives sequence designs directly from the agent and constructs the corresponding enzyme variants through an automated DNA assembly workflow^46^. The resulting genes are expressed in a cell-free protein expression system and characterized using fluorescence-based enzyme assays in a multimode plate reader. All DNA fragments and reagents are stored on-deck, allowing the system to progress from sequence design to functional measurement in approximately 10 hours (Fig. 2c). Across the entire autonomous engineering campaign, the laboratory successfully assembled 136 of 150 designed variants (91%) and generated high-quality assay measurements for 132 of those assemblies (97%), demonstrating robust operation over extended multi-week campaigns (Fig. S2).

The laboratory is tightly integrated with the computational agent through a common software infrastructure that coordinates experiment execution, data management, and model training (Fig. 2b). The laboratory control computer and GPU server hosting the agent are physically separated but communicate automatically over the network, with experimental measurements returned for model updating and newly proposed variants sent to the laboratory for construction and testing. Robust error handling allows the system to continue operating despite occasional assembly failures or assay-quality issues. Together, these capabilities enable continuous generation of experimental feedback for autonomous learning.

### Autonomous multi-agent engineering of glycoside hydrolase specificity

We next challenged the autonomous agents to engineer substrate specificity in glycoside hydrolases (GHs). GHs are among the most abundant and important enzyme families in nature, playing central roles in biomass degradation, carbon cycling, and biotechnology^47^. We focused on Family 1 glycoside hydrolases (GH1s)^48^, which have evolved high activity and specificity toward glucose-containing substrates, reflecting the central importance of glucose metabolism across biology. Reprogramming this preference toward other sugars is therefore a challenging protein engineering problem, particularly because many sugars differ from glucose by only subtle structural features. As a result, engineering GH1 substrate specificity provides a stringent test of whether autonomous experimental interaction can discover enzyme functions that are difficult to predict or engineer.

We formulated autonomous experimental interaction as a coordinated multi-agent problem. Three autonomous agents were assigned complementary engineering objectives: Agent G prioritized glucose specificity, Agent X prioritized xylose specificity, and Agent M prioritized mannose specificity. Each agent independently proposed variants for experimental evaluation using its own acquisition policy, while all observations were incorporated into a shared experimental memory and common learning model. As a result, observations made under one objective immediately informed subsequent decisions across the system. This multi-agent framework provides a path toward scaling autonomous experimental interaction by enabling multiple objectives to be explored simultaneously while accumulating knowledge through a shared experimental experience (Fig. 3a). We hypothesized that optimizing activity alone would favor promiscuous enzymes, whereas optimizing specificity alone would favor weakly active enzymes. We therefore tasked each agent with maximizing an objective that combines both properties: substrate specificity, defined as the fraction of total activity toward the target substrate, multiplied by activity toward that substrate.

**Figure 3.**
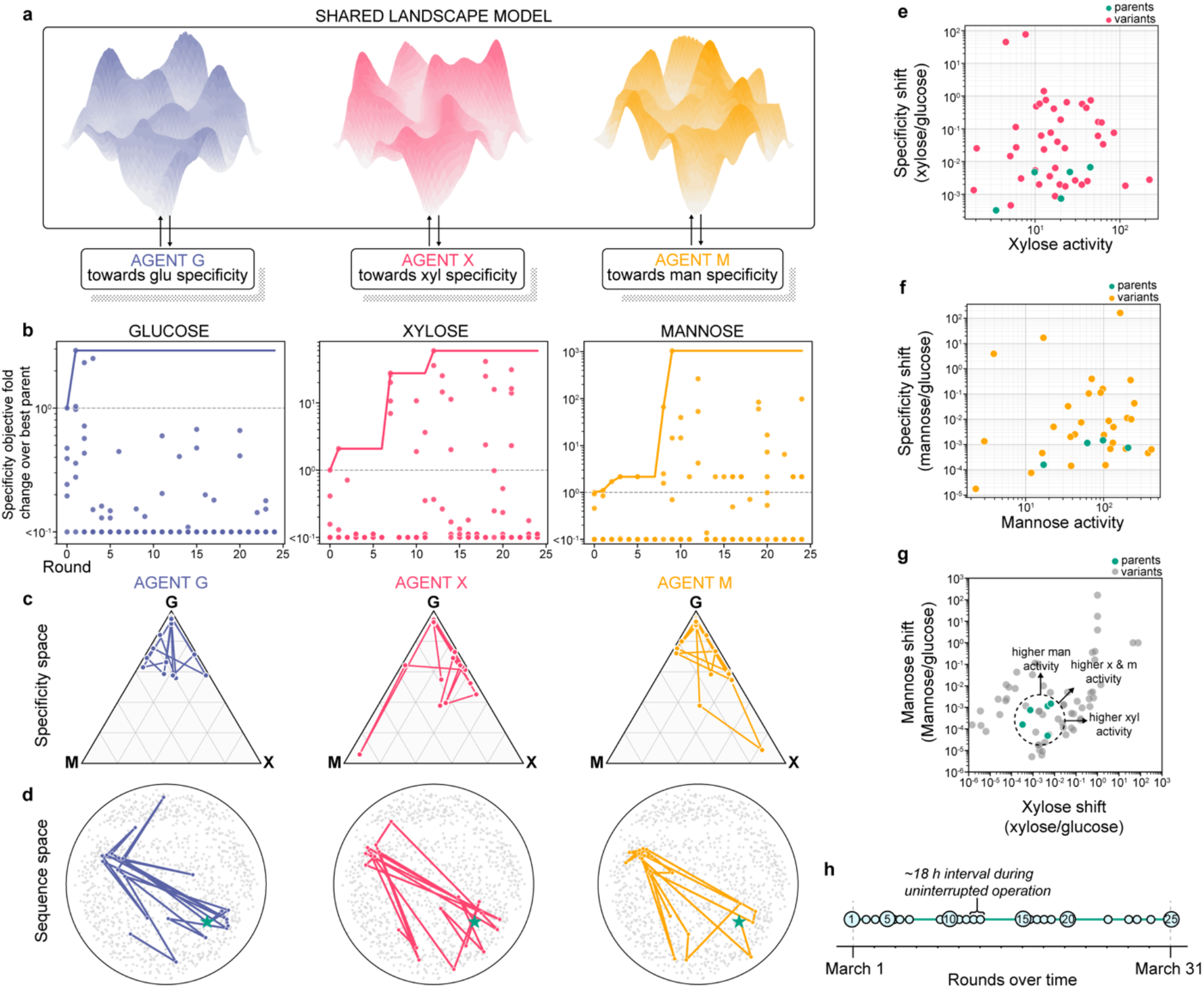
Multi-agent exploration of glycoside hydrolase specificity. (a) Three autonomous agents pursue distinct substrate objectives while learning from a shared experimental memory and model of the sequence–function landscape. (b) Specificity objective trajectories over the autonomous experimental campaign, showing measured values (circles) and the cumulative best value (solid line) for each substrate. (c) Search trajectories of the three agents in ternary specificity space. (d) Search trajectories of the three agents in dimensionally reduced (MDS) sequence space. (e) Xylose specificity shift (ratio of xylose to glucose activity) versus xylose activity for all characterized variants, showing the discovery of variants with increased xylose specificity and activity. (f) Corresponding specificity– activity relationship for mannose. (g) Xylose versus mannose specificity shifts for all characterized variants, showing exploration beyond the narrow substrate preferences of the parent enzymes. (h) Timeline of the autonomous campaign, comprising 25 rounds of experimental interaction over approximately one month.

The campaign was initialized using six diverse naturally occurring GH1 enzymes. We defined a combinatorial sequence space by recombining eight sequence fragments from these six parent enzymes, generating approximately 1.7 million possible chimeric variants while preserving the overall GH1 architecture (Fig. S3). At each round, the agents proposed new enzyme variants, the autonomous laboratory constructed and characterized them, and the resulting measurements were incorporated into the shared model to guide subsequent decisions. The system operated continuously for approximately one month, completing 25 rounds of autonomous experimentation (Fig. 3h). During this period, the platform autonomously designed enzyme variants, constructed and tested them in the laboratory, analyzed the resulting data, updated its internal models, and selected subsequent experiments without human intervention.

Over the month-long experiment, the autonomous learning system explored a broad range of substrate preferences and progressively discovered variants with improved specificity across all three objectives. Improvements toward glucose were modest, increasing 2.9-fold over the best parent enzyme, reflecting the already strong native preference of these enzymes for glucosyl substrates. In contrast, substantial gains were achieved toward the non-native xylose and mannose objectives, with specificity increasing approximately 60-fold and 1000-fold, respectively (Fig. 3b). However, these fold changes can become large and uncertain when the specificity of the reference parent approaches the assay detection limit. The individual search trajectories towards each substrate also differed substantially across objectives: Specificity toward xylose increased gradually throughout the campaign, with significant improvements emerging over multiple rounds of iterative learning. In contrast, specificity towards mannose improved through a smaller number of significant discrete jumps, including large gains in rounds 8 and 9 that persisted through subsequent rounds. These differences likely reflect both the underlying chemistry of the substrates and the information available to the learning system. The initial GH1 enzymes displayed substantially higher activity toward xylose than toward mannose, with many variants falling near the detection limit on the latter. As a result, exploration toward xylose generated richer and more continuous experimental feedback, whereas improvements toward mannose depended on identifying relatively rare variants with measurable activity. Despite operating under these distinct regimes, the autonomous system successfully engineered specificity toward both substrates, demonstrating its ability to learn from experimental feedback across objectives with markedly different fitness landscapes and signal strengths (Fig. 3d,e). Collectively, the three agents discovered variants spanning substrate preferences well beyond the narrow specificity range of the parent enzymes (Fig. 3g).

To verify these discoveries, we manually repeated experiments on a set of top-performing variants alongside the initial parent enzymes. The results confirmed that engineered variants displayed significantly increased specificity toward their target substrates. Across the variants selected by each agent, Agent G preferentially identified glucose-specific variants, whereas Agents X and M were enriched for variants with increased specificity toward xylose and mannose, respectively (Fig. S4).

### Learning substrate specificity through autonomous experimental interaction

To rigorously quantify the specificity changes identified during autonomous exploration, we characterized a panel of engineered variants and the six parent GH1 enzymes using purified-protein kinetic measurements across all three substrates. Because many enzymes exhibited substrate inhibition, we determined catalytic efficiency (*k*_*cat*_*/K*_*m*_) from the initial low-substrate slope of the velocity curve. As expected, all parent enzymes strongly preferred glucose, with catalytic efficiencies 460-to 1118-fold lower toward xylose and 1300- to 7000-fold lower toward mannose (Fig. 4a). Several engineered variants exhibited substantially altered specificity toward the non-native substrates, including X1, M7, X8, and M21, which exceeded the specificity of their constituent parent enzymes (Fig. 4b). In the opposite direction, two variants discovered by Agent G, G1 and G9, exhibited complete glucose specificity, with no detectable activity toward xylose or mannose. Collectively, the three agents also discovered variants that combined increased activity with increased specificity toward both non-native sugars (Fig. 4c,d). These results confirm that autonomous exploration produced substantial changes in catalytic specificity that persisted under purified-enzyme characterization.

**Figure 4.**
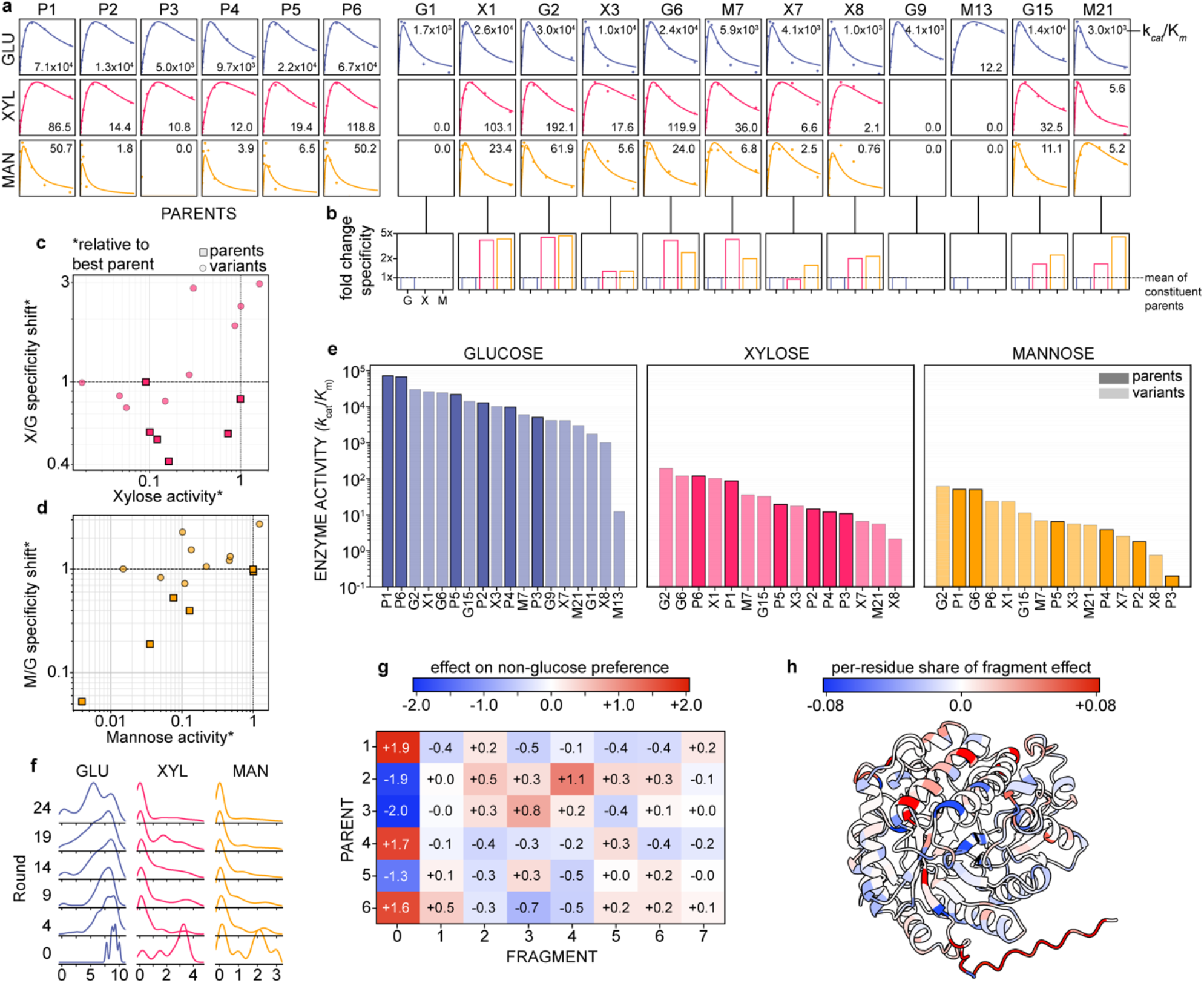
Learning glycoside hydrolase substrate specificity through experimental interaction. (a) Purified-protein kinetic characterization of the six parent enzymes and twelve engineered variants across glucose, xylose, and mannose, with catalytic efficiencies (k_*cat*_/K_*m*_) shown for each enzyme–substrate pair. (b) Fold-change in substrate specificity of each engineered variant relative to its constituent parent enzymes. For each variant and substrate, specificity is normalized to the mean specificity of the parent enzymes contributing sequence fragments to that variant. Colored circles show the corresponding mean across all six parent enzymes. (c) Xylose specificity shift versus xylose catalytic efficiency, highlighting variants that combine increased xylose activity and specificity. (d) Corresponding activity–specificity relationship for mannose. (e) Catalytic efficiencies of kinetically characterized enzymes across the three substrates. (f) Predicted activities of the same 1,000 randomly sampled chimeras using model checkpoints from successive rounds of autonomous experimentation, revealing how the model’s inferred sequence–function landscape changed with accumulated experimental experience. (g) Effect of each parental fragment on predicted non-glucose preference, quantified as [(Xyl + Man)/2 − Glu], relative to the mean across the same 1,000 random chimeras using the final model checkpoint. (h) Residue-level decomposition of these fragment effects mapped onto the GH1 structure, identifying sequence positions associated with predicted substrate preference.

Analysis of the engineered variants revealed a shared relationship between the two non-native specificity objectives. Increased xylose and mannose specificity were strongly correlated, driven largely by decreased glucose activity rather than parallel gains in activity toward both non-native substrates. Consistent with this coupling, variants selected by Agent X and Agent M occupied similar regions of sequence space and shared related sequence compositions (Fig. S5). Together, these observations suggest that many sequence changes discovered during autonomous exploration shift substrate preference primarily by disrupting the strong native glucose preference of GH1 enzymes, thereby benefiting both non-native specificity objectives. Because these changes were informative for both objectives, experimental observations generated under one objective could also inform the other.

We next interrogated the learned model to understand what knowledge the agent had acquired through experimental interaction. We first examined how its predictions evolved over successive rounds of the campaign. We scored the same 1,000 randomly sampled chimeras using model checkpoints from successive rounds, providing a consistent probe of how the inferred landscape changed as the agent accumulated experimental experience. Initially, having observed only the six parent enzymes, the model predicted that most sequences across the combinatorial space would retain relatively high activity. As experimental observations accumulated, this picture changed substantially (Fig. 4f). By the final round, the model predicted a broad distribution of glucose activities, including populations of inactive and weakly active variants, while predicted activities toward xylose and mannose were strongly right-skewed, with high activity confined to progressively rarer regions of sequence space. Thus, autonomous experimentation progressively revealed a landscape very different from that suggested by the parent enzymes alone: glucose activity was broadly accessible, xylose activity was less common, and mannose activity was comparatively rare.

We next asked what sequence determinants of substrate preference the agent had learned through experimental interaction. Using the final model checkpoint, we quantified predicted non-glucose preference for the same 1,000 random chimeras as the average predicted activity toward xylose and mannose relative to glucose, [(Xyl + Man)/2 − Glu]. We then measured how the presence of each parental fragment shifted this predicted preference relative to the average across all chimeras (Fig. 4g). This analysis revealed that substrate preference was strongly influenced by a small number of sequence regions, with particularly large effects associated with the first fragment. In contrast, fragments containing the two catalytic glutamate residues had relatively little influence on predicted non-glucose preference, suggesting that substrate specificity is distributed across the protein and strongly influenced by regions outside the catalytic center^49^.

We further decomposed these fragment-level effects to identify individual residues associated with predicted substrate preference and mapped their contributions onto the GH1 structure (Fig. 4h). The strongest effects were distributed throughout the protein, although several occurred near the central substrate-binding pocket of the GH1 α/β barrel. This pattern suggests that substrate preference emerges from distributed sequence changes that may influence substrate recognition both directly within the substrate-binding region and indirectly through the surrounding protein structure. These model-derived associations provide testable hypotheses for the molecular determinants of GH1 substrate specificity.

Beyond substrate specificity, autonomous exploration revealed an unexpected sequence determinant associated with protein expression. Three of the highest-activity variants identified during the month-long experiment shared the same sequence fragment, P2f0 (fragment 0 from parent 2), despite arising from distinct search trajectories. Follow-up characterization showed that these variants exhibited substantially increased protein expression, with one producing expression-associated cytotoxicity. Because expression was never an explicit optimization objective, this relationship emerged from observations accumulated during autonomous exploration. This finding illustrates how experimental interaction can reveal biological relationships beyond those explicitly targeted by the agent^50^.

## Discussion

AI-based protein structure prediction, protein language models, and generative design methods have transformed biology by learning from vast collections of naturally evolved sequences and experimentally characterized proteins. Yet these approaches remain largely observational, learning from existing data rather than generating new knowledge through action and experience. Autonomous experimental interaction provides a complementary paradigm in which AI can perturb biological systems, observe the consequences, and iteratively refine its understanding through experimental feedback. Here, we demonstrate this approach by coupling a generative protein language model with a fully automated laboratory, enabling an autonomous agent to design, construct, and characterize enzyme variants and learn from the resulting experimental feedback. Applied to glycoside hydrolases, autonomous exploration over approximately one month produced variants with substantially altered substrate specificity toward non-native sugars. More broadly, these results establish experimental interaction as a framework for AI to learn biological function through experience.

The autonomous exploration also provides insight into how the structure of the fitness landscape and experimental feedback shape interactive learning. Specificity toward xylose improved gradually over many rounds, whereas mannose specificity emerged through a smaller number of large jumps. These distinct trajectories likely reflect differences in both the underlying fitness landscapes and the information available to the learning system. Initial GH1 activity toward xylose was substantially higher than toward mannose, providing richer and more continuous experimental feedback and a more accessible path toward improved function, while mannose activity frequently approached the detection limit. More generally, these results suggest that the efficiency of autonomous exploration depends both on the structure of the underlying fitness landscape and on the quality and dynamic range of the measurements used to navigate it.

Beyond engineering altered enzyme specificity, autonomous exploration generated new insight into the underlying sequence–function landscape. Kinetic characterization revealed that specificity differences were driven primarily by changes in catalytic turnover rather than substrate binding: across the parent enzymes, differences in *k*_*cat*_ were substantially larger than differences in *K*_*m*_, suggesting that discrimination among these closely related sugars occurs largely during catalysis rather than initial substrate recognition. Analysis of the learned model further identified distributed sequence determinants of substrate preference, including contributions from regions outside the catalytic center, providing testable hypotheses for how GH1 specificity is encoded. Finally, several high-activity variants independently converged on a common parental fragment associated with markedly increased protein expression, despite expression never being an explicit optimization objective. Together, these observations illustrate how autonomous experimental interaction can do more than search for improved function: the resulting experience can reveal relationships within the biological system that were not specified in advance.

The multi-agent experiment illustrates how shared experimental experience can increase the information available to individual learners. Each agent independently selected variants according to its own substrate objective, but every resulting measurement was incorporated into the common training data used by all three agents. Thus, while each agent was responsible for only approximately one third of the experiments, it learned from the complete set of 132 successfully characterized variants. This sharing was particularly relevant for the xylose and mannose objectives, which proved to be coupled through common changes in substrate preference: observations generated while pursuing one objective could therefore provide information relevant to the other. Indeed, some of the strongest variants for each substrate were discovered by agents pursuing a different objective. Although we did not directly compare shared and isolated agents, this architecture provides a simple mechanism for improving sample efficiency by allowing each experiment to inform multiple learning objectives rather than benefiting only the agent that selected it.

This principle provides a path toward scaling experimental learning. Unlike purely computational learning, the rate of experimental learning is ultimately constrained by the rate at which physical experiments can be performed. Increasing numbers of autonomous agents and laboratories could overcome this constraint through parallel experimentation coupled with shared learning. Future agents could also coordinate more explicitly, dividing scientific problems into complementary subproblems, exploring different regions of sequence space, testing competing mechanistic hypotheses, or pursuing orthogonal functional objectives. These agents need not operate within a single laboratory: distributed networks of autonomous laboratories could generate experimental experience in parallel while contributing observations to shared models. In this way, scaling the number and diversity of experimental agents could expand both the rate and scope of biological learning.

More broadly, autonomous experimental interaction represents a shift from biological AI that learns primarily from existing observations toward AI that can acquire knowledge through experience. Future agents could continuously perturb biological systems, observe the consequences of their actions, and use those observations to refine their understanding of biological function. Just as large-scale observational datasets enabled the recent revolution in AI for biology, large-scale autonomous experimentation may enable a new era of biological discovery driven by interaction.

## Methods

### Automated laboratory

The automated laboratory consists of a Dynamic Devices Lynx LM730i liquid handler, Analytik Jena Biometra TRobot II thermal cycler, Agilent BioTek Synergy H1 plate reader, and custom-built automated refrigerator arranged around a central Peak Robotics KX2-750 robotic arm. The robotic arm transfers labware and reagents between instruments, enabling automated execution of the complete protein engineering workflow. Laboratory instruments are controlled by a dedicated computer that communicates over the network with the computational agent running on a physically separate GPU server. Sequence designs are transferred from the GPU server to the laboratory control computer via HTTP, while experimental measurements are returned to the GPU server via secure file transfer protocol (SFTP). This architecture allows computational model training and experimental execution to operate independently while automatically exchanging designs and measurements between successive rounds.

### Automated protein production and characterization

We used a chimeric fragment assembly procedure to generate a large combinatorial library of protein variants. Six homologous Family 1 glycoside hydrolase (GH1) sequences from the soil-dwelling bacterial genus *Streptomyces* served as the parent enzymes. Each parent sequence was divided into eight fragments at minimally disruptive sites using the SCHEMA recombination algorithm^51,52^. Fragments at each position could be drawn from any of the six parents, yielding a combinatorial sequence space of 6^8^ = 1,679,616 possible chimeric variants.

The laboratory was initialized with all reagents required for autonomous protein construction and characterization in the on-deck refrigerator. These included the 48 parental DNA fragments stored on plasmids (320 ng/μL), 2× Golden Gate Assembly Master Mix prepared from the NEBridge Golden Gate Assembly Kit (E1601L), NEB Phusion Hot-Start Flex DNA Polymerase (M0535L), forward and reverse PCR primers (2 μM), EvaGreen dsDNA dye, and reagents from the Bioneer AccuRapid Cell Free Protein Expression Kit (K-7260). The cell-free expression mixture contained AccuRapid *E. coli* extract supplemented with 40 μM fluorescein as an internal standard. Three fluorogenic substrate solutions contained 138.9 μM 4-methylumbelliferyl-β-D-glucopyranoside, 4-methylumbelliferyl-β-D-xylopyranoside, or 4-methylumbelliferyl-β-D-mannopyranoside in 0.56% (v/v) DMSO, 11 mM phosphate, and 56 mM NaCl, pH 7.0.

At the beginning of each experimental round, the laboratory assembled agent-selected combinations of DNA fragments by Golden Gate assembly. DNA fragments were combined in a thin-walled PCR plate, mixed 1:1 with 2× Golden Gate Master Mix, and subjected to 60 cycles alternating between 37 °C and 16 °C for 1 min each. Assembly products were PCR-amplified using Phusion polymerase (30 s at 98 °C; 30 cycles of 30 s at 98 °C, 30 s at 59 °C, and 60 s at 72 °C; final extension for 5 min at 72 °C). PCR products were diluted and quantified using EvaGreen fluorescence against a dsDNA standard curve. Assemblies exceeding 43 ng/μL were advanced to protein expression, whereas failed assemblies were returned to a retry pool available to the computational agent in subsequent rounds. Successful assemblies were expressed for 3 h at 30 °C using the AccuRapid cell-free expression system, with a no-DNA reaction included as a negative control. Each expression reaction was then assayed in triplicate against glucose-, xylose-, and mannose-derived fluorogenic substrates in 96-well plates. Fluorescein fluorescence (excitation 487 nm, emission 528 nm) was measured as an internal expression standard, followed by enzyme activity measurements (excitation 325 nm, emission 450 nm) every 5 min for 1 h.

Upon completion, the assay plate was discarded and fresh labware was loaded automatically for the next experimental round.

### Automated enzymatic characterization

Rather than acquiring full substrate-concentration series for every variant, we used the initial rate (*V*_*0*_) at a single substrate concentration as a proxy for activity. The parent enzymes exhibited substrate inhibition in cell-free lysate, with rates increasing to a maximum and then declining at higher substrate concentrations. We therefore performed assays near the rate-maximizing concentration (125 μM; Fig. S6). At fixed substrate concentration and enzyme level, *V*_*0*_ scales with *V*_*max*_ and therefore *k*_*cat*_, with additional dependence on *K*_*m*_ and *K*_*i*_. Purified-enzyme kinetic measurements showed that differences among these enzymes were driven primarily by *k*_*cat*_ rather than *K*_*m*_ (Supplementary Data), supporting the use of *V*_*0*_ as a *k*_*cat*_-weighted activity proxy. The single-point assay therefore does not directly measure *k*_*cat*_/*K*_*m*_, but captures catalytic differences that contribute strongly to substrate specificity in this system. Substrate turnover was monitored by fluorescence (excitation 325 nm, emission 450 nm) every 5 min for 1 h (13 timepoints). Each enzyme was assayed in triplicate against glucose, xylose, and mannose, with no-enzyme controls included for each substrate.

For each well, reaction rate was estimated from the slope of fluorescence over time. For most wells, the rate was obtained by ordinary least-squares regression over the full time course. For rapidly turning-over reactions (full-trace slope >30 RFU/s), substrate depletion or detector saturation could introduce curvature and bias the full-trace estimate. These reactions were additionally fit using a continuous piecewise-linear model, *y*(*t*) = *a* · *t* + *b* + *c* · max(0, *t* – *t*_*pb*_), where *a* is the initial slope and *t*_*bp*_ is the breakpoint. The breakpoint was selected by exhaustive search over candidate timepoints, requiring at least three points per segment, to minimize the residual sum of squares. The four-parameter piecewise model was used only when favored over the two-parameter linear model by the Bayesian information criterion, 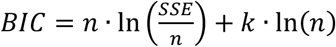, where *k* is the number of model parameters; otherwise, the full-trace linear slope was retained. Fluorescence measurements exceeding the linear detection range were excluded by truncating the time course at the first saturated reading before fitting.

Per-well rates were corrected for variation in cell-free expression yield using the co-measured fluorescein endpoint signal (excitation 487 nm, emission 528 nm), normalized within each enzyme–substrate replicate group. Background turnover was removed by subtracting the median no-enzyme control rate from the median replicate rate for each enzyme–substrate combination. Negative values were set to zero, and the resulting rates (mRFU/s) were transformed as ln(1 + *x*) to produce the activity phenotype used for model training.

### Model training and variant generation

Our surrogate model for protein function prediction is ProteinNPT, a variant of a non-parametric transformer. After characterization, sequences are sent with Secure File Transfer Protocol (SFTP) to the GPU server, where they are added to the training database. A thorough explanation of model training can be found in the ProteinNPT manuscript. However, briefly: The corresponding zero-shot scores for each sequence are computed using the autoregressive protein language model Tranception, which are appended as auxiliary labels and passed as input but not masked during training (and thus do not contribute to the loss). Additionally, a binary activity label is appended, which is passed as a normal input label that is masked and whose loss is backpropagated during training. These data are embedded into ProteinNPT: protein sequences using ESM2-650M, and the associated labels, including activity toward each of the three substrates, zero-shot scores, and a binary label indicating whether the variant exhibited detectable activity, using a learned linear transformation. The concatenated dataset is passed through many successive ProteinNPT layers, where 15% of input sequence tokens and all label tokens are masked. The algorithm applies axial attention along the rows and columns to model the dependencies between all tokens in the input dataset. It does so by optimizing the loss:

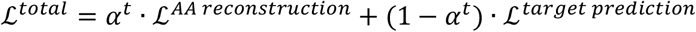

where ℒ^*AA reconstryuction*^ is a cross-entropy loss over amino acid tokens and ℒ^*target prediction*^ is a mean-squared error (MSE) loss over activity target predictions. αt is a weighting term that is progressively annealed to focus training on target prediction of activity labels. All relevant hyperparameters can be found in Supplementary table 1.

At variant generation, the behavior of the three agents (each having the objective to optimize specificity on their respective substrate) diverges: each of the three agents samples, scores, and acquires sequences independently while sharing the same training data and trained model.

At variant generation, each agent begins by first defining its specificity objective function. We hypothesized that if we programmed each agent to optimize pure enzyme activity on its corresponding substrate, they would be biased to produce promiscuous enzymes. Similarly, if we programmed each to optimize pure specificity, they would be biased to produce lowly active enzymes. We use our specificity objective function to generate enzymes that display both activity and specificity. Given *A*_*glu*_ as the activity phenotype toward glucose generated from the automated laboratory, we define the glucose specificity objective as:

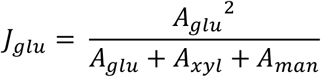

This objective is the product of glucose activity and glucose specificity, *A*_*glu*_/(*A*_*glu*_ + *A*_*xyl*_ + *A*_*man*_), and was chosen to favor variants with both high activity and high specificity toward the target substrate. Objectives for xylose and mannose were calculated similarly.

Candidate variants were generated such that each differed by a single fragment from any previously characterized sequence in the accumulated experimental dataset. To bias generation toward high-performing regions of sequence space, previously characterized sequences were ranked according to specificity for the agent’s substrate objective, and the top quartile was used to condition fragment sampling. The activity labels of these conditioning sequences were set to values corresponding to the maximum observed specificity in the training dataset, providing the conditioning signal for sampling. Once the conditioning pool was formed, sequences and their segmented representations were used to generate candidate variants in batches until the desired sample count was reached. Each iteration draws a random permutation over the fragment indices for every parent, so that masking proceeds in a different order across sequences. For a given fragment position *k*, we replace the corresponding residues in every parent with the special token “<mask>“ and concatenate the segments to obtain full-length masked sequences. These masked sequences, together with mutation strings relative to the wild type, are passed into the trained ProteinNPT model. The model restores the amino-acid embeddings, produces logits for every position, and we apply a temperature-scaled softmax (default temperature *T* = 1) to obtain log-probabilities over the 20 amino acids.

We then isolate the log-probabilities covering fragment k for each sequence and feed them to a chunk sampler that evaluates all candidate fragments in the precomputed library. The sampler aggregates token-level likelihoods over each fragment, renormalizes them, and returns a replacement fragment *s*^*k*^. The sampled fragment is spliced back into the sequence. Cycling through all *K* fragments across all *B* parents yields a batch of offspring. Outer iterations repeat until the targeted number of mutants (default = 1000) is produced. We then filter the generated set to remove duplicates and any sequence already present in the acquisition database.

### Acquisition

Predictive uncertainty is estimated via Monte Carlo dropout^53^ with five stochastic forward passes through ProteinNPT with no masking, providing per-target means μ and standard deviations σ for each candidate sampled in the batch. Acquisition scoring follows an Upper Confidence Bound strategy. For substrate *i* the score is:

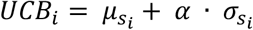

where 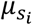 and 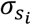 denote the posterior mean and predictive uncertainty of specificity toward substrate *i*. The hyperparameter α (default *α* = 1.0) governs the trade-off between exploration and exploitation; low values of α bias the algorithm toward exploitation, relying heavily on current knowledge at the expense of discovering new optima. Conversely, excessively high values of α drive the algorithm toward exploration, prioritizing highly uncertain regions of the landscape that often yield non-functional variants. Setting *α* = 1.0 places equal weight on the predicted mean and the predictive uncertainty, ensuring the algorithm pursues high-potential novel variants without being misled by excessive uncertainty. The sequence to acquire for lab characterization is then:

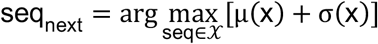

Each of the three agents identifies two sequences to query the lab for experimental characterization.

### In-silico sequence space exploration benchmarks

We benchmarked our method on four diverse protein DMS datasets. Each simulated engineering campaign was initialized with 10 sequences drawn uniformly at random from the dataset and run for 20 rounds, with 10 sequences selected per round (200 oracle queries, corresponding to 0.1–0.5% of each available landscape). At each round, ProteinNPT was retrained on all labeled sequences. Candidate variants were then generated around up to 30 sequences from the top quartile of the currently labeled set. For each round, a Hamming distance n was sampled from a Poisson distribution (λ = 1), with *n* = 0 replaced by *n* = 1. All variants exactly n mutations from the selected sequences were enumerated, and the resulting candidate pool was randomly subsampled to 1,000 sequences when necessary. Candidates were scored using ProteinNPT, and the next 10 sequences were selected by UCB acquisition (*α* = 1.0) using MC-dropout uncertainty. Experimental measurements were simulated by revealing the corresponding fitness values from the DMS dataset.

To isolate the contributions of the model representation and acquisition strategy, we compared five search strategies, each evaluated across five random seeds per protein: (1) ProteinNPT with UCB acquisition; (2) ProteinNPT with greedy acquisition (*α* = 0); (3) a 10-member one-hot MLP ensemble with UCB acquisition, using the ensemble mean as predicted fitness and ensemble standard deviation as uncertainty; (4) the same MLP ensemble with greedy acquisition; and (5) uniform random selection. Comparing ProteinNPT with the MLP ensemble isolates the contribution of pretrained protein representations relative to a supervised model trained from scratch on the same labels, while comparing UCB with greedy acquisition isolates the contribution of uncertainty-guided exploration. Uniform random selection provides a no-model baseline. Search trajectories are reported as the per-round median across the five random seeds, with final values from individual runs shown separately to illustrate run-to-run variability (Fig. S1).

### Quantifying landscape epistasis

For each protein we characterize its top 1% fitness region (the regime in which the searches converge) along two complementary axes: how far local fitness departs from additivity, and what geometric (epistatic) type that departure takes. We first decompose fitness into additive and epistatic components. An additive model encodes each variant as a binary vector over its constituent single–amino-acid substitutions and predicts fitness by ridge regression (*α* = 1.0); its coefficient of determination (additive R^2^) is the fraction of regional fitness variance captured by context-independent per-mutation effects. We then fit a global-(nonspecific-) epistasis model by passing the additive prediction through a single monotonic transform estimated by isotonic regression (monotone R^2^). Because the transform is monotonic it preserves the rank order of the additive score and so introduces no new local optima or sign reversals; the gain of monotone R^2^ over additive R^2^ therefore measures navigable, nonspecific epistasis (saturation and diminishing-returns effects), while the residual variance (1 − monotone R^2^) reflects specific, genotype-dependent epistasis together with measurement noise. Both models are refit within the region, since regularized regional fits represent the peak more faithfully than extrapolations of a globally fit model. We resolve the geometric type of local epistasis on the genotype graph. For every pair of mutations *a, b* sharing a measured background *R* for which all four corners *R, R* + *a, R* + *b, R* + *a* + *b* are assayed. We compute the interaction term *ε* = *f*(*R* + *a* + *b*) − *f*(*R* + *a*) − *f*(*R* + *b*) + *f*(*R*) (wild-type–anchored pairs use the measured wild-type fitness as baseline). Each face is classified from the sign behaviour of its two marginal effects: when neither mutation’s effect changes sign between backgrounds the interaction is *magnitude* epistasis, when exactly one changes sign it is *sign* epistasis, and when both change sign it is *reciprocal sign* epistasis.

To avoid attributing measurement noise to epistasis, we classify each face against a per-face noise floor rather than against zero. Because *ε* is a linear combination of four measurements at mutation orders (|*R*|, |*R*| + 1, |*R*| + 2), its noise variance is the sum of the four corner variances *σ*^2^_ε;_(*face*) = *σ*^2^_{|*R*|}_ + 2*σ*_{|*R*|+1}_ + *σ*^2^_{|*R*|+2}_. The single-mutant measurement noise *σ*_/1%2)-_ is taken from each study’s replicate reliability *r* via the relation *σ*^2^_*single*_ = (1 − *r*) · *var*(*singles*) (the “r-trick”) for GB1 (*r* = 0.996) and Ube4b (*r* = 0.89), and directly from the reported wild-type or synonymous-variant control noise for avGFP and Pab1, where library-wide replicate correlations were not available (equivalent *r* = 0.98 and 0.91). Higher-order corner noise is set equal to *σ*_*single*_ except where a study reports it separately: GB1’s double-mutant measurements are far noisier than its singles owing to lower read depth rather than greater signal variance (the variance of doubles and singles is comparable), so we use the directly reported double-mutant noise (≈0.52). A face is called non-epistatic when |*ε*| ≤ *τ*, and a marginal sign flip is counted only when both the alone- and in-background effects exceed *τ* in magnitude, with the threshold fixed at one face standard deviation (*τ* = *σ*_*ε*_). For each protein we report monotone R^2^, and the noise-floored partition of its top-1% faces into non-epistatic, magnitude, sign, and reciprocal sign classes.

### Lysate replication experiments

To validate a set of top-performing sequences from the automated enzyme engineering campaign plus the six parents, we manually replicate the enzymatic activity assays across all three substrates in identical lysate and assay conditions. 3.125 ng/μL of plasmid DNA containing each enzyme variant was added in 100 μL total cell-free expression volume and incubated at 30 °C for 3 hours. The resultant expression was pipetted in 10 μL volumes across 3 replicates of each of the 3 substrates. A corresponding set of negative controls, containing water (0 ng/μL DNA) added to the cell-free expression, was pipetted in similar 10 μL volumes. 90 μL of substrate buffer was mixed with each, and the 96-well plate was read at ex: 325 nm/em: 450 nm every 5 minutes for 1 hour. After applying correction from the corresponding negative controls by per-time-point subtraction, the initial slopes were extracted from the time courses, compared to a standard curve of 4MU fluorescence, and used to represent each variant’s activity as a rate of 4MU production·s^-1^. We represent enzyme specificity in Fig. S3 as a fraction of total activity, e.g. for glucose (and similarly calculated for the other substrates):

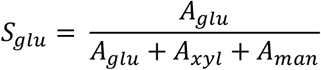

### Enzyme expression and purification

Sequences encoding the six parent enzymes and selected engineered variants were ordered from Twist Bioscience in pET-blank plasmids. Plasmids were transformed into *E. coli* BL21(DE3) and plated on LB agar containing 50 μg/mL carbenicillin. Following overnight growth at 37 °C, individual colonies were used to inoculate 5-mL LB starter cultures containing 50 μg/mL carbenicillin and grown overnight at 37 °C with shaking at 250 rpm. Starter cultures were diluted 100-fold into 50 mL LB containing 50 μg/mL carbenicillin and grown at 37 °C with shaking at 250 rpm to an OD600 of 0.4–0.6. Protein expression was induced with 1 mM isopropyl β-D-1-thiogalactopyranoside (IPTG), and cultures were incubated overnight at 16 °C with shaking. Cells were harvested by centrifugation at 4,000 × g for 20 min.

Cell pellets were resuspended in lysis buffer (25 mM Tris, 400 mM NaCl, 10 mM imidazole, 10% v/v glycerol, pH 7.5), incubated on ice for 30 min, and lysed by sonication. Lysates were clarified by centrifugation at 15,000 × g for 15 min and loaded onto gravity columns containing Ni-NTA resin (Thermo Scientific HisPur). Columns were washed with 20 mL wash buffer (25 mM Tris, 400 mM NaCl, 20 mM imidazole, 10% v/v glycerol, pH 7.5), and proteins were eluted with 5 mL elution buffer (25 mM Tris, 400 mM NaCl, 250 mM imidazole, 10% v/v glycerol, pH 7.5). The enzyme samples were analyzed via SDS-PAGE to quantify purity (Fig. S7). The samples were then buffer exchanged and concentrated to approximately 1 mg/mL in storage buffer (25 mM Tris, 100 mM NaCl, 10% v/v glycerol, pH 7.5) using Amicon centrifugal filters. Protein concentrations were determined by bicinchoninic acid (BCA) assay. Purified proteins were flash-frozen in liquid nitrogen and stored at −80 °C until use.

### Michaelis-Menten kinetics

Purified enzymes were assayed against the three fluorogenic substrates 4-methylumbelliferyl-β-D-glucopyranoside, -xylopyranoside, and -mannopyranoside (4MU-glu, 4MU-xyl, 4MU-man) in 10 mM sodium phosphate, 50 mM NaCl, 0.5% v/v DMSO, pH 7.0, in a 96-well microplate at 30 °C. Each enzyme–substrate combination was run in triplicate over an eight-point two-fold serial dilution of substrate (ranges given below). Reaction progress was followed by monitoring 4MU fluorescence on a microplate reader (excitation 325 nm, emission 450 nm). A matched plate of no-enzyme negative controls was run in identical conditions for each substrate and subtracted per timepoint at the fluorescence level prior to slope estimation. For each concentration well, v_0_was obtained by ordinary least-squares regression over the linear region of the background-subtracted progress curve (windows specified below), converting the resulting fluorescence slope to 4MU product (µM·s^−1^) using a 4MU standard curve measured on the same instrument; triplicate v_0_values were averaged at each substrate concentration. Steady-state parameters were then estimated by fitting the substrate-inhibition (Haldane) model:

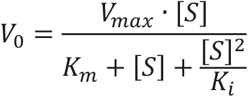

to v_0_versus [S] by non-linear least squares (scipy.optimize.curve_fit) over a broad grid of starting guesses; *k*_cat_ was computed as *V*_max_/[E]. Because *K*_m_ and *V*_max_ are statistically correlated in the substrate-inhibition fit, *k*_cat_/*K*_m_ is instead estimated model-independently from the low-[S] regime, where the model reduces to v ≈ (*V*_max_/*K*_m_)·[S]: an ordinary least-squares fit of v_0_against [S] across the three lowest substrate concentrations recovers *V*_max_/K_m as the slope, which divided by [E] gives the reported *k*_cat_/*K*_m_. Because the Haldane fit does not always converge (and in this case, left several enzymes’ kinetic parameters anti-correlated and poorly constrained), we report *k*_*cat*_*/K*_*M*_ from the initial slope of *V*_0_ vs. [S] at low substrate concentrations as a model-free specificity measurement for every enzyme-substrate pair.

*Parent enzymes (P1–P6)*. Parents were assayed at 10 nM enzyme on glucose and 100 nM enzyme on xylose and mannose. The glucose substrate dilution series ran from 500 µM down to 3.9 µM and was monitored every 2 min for 30 min (16 timepoints) at a gain setting of 80; the xylose and mannose dilution series ran from 1000 µM down to 7.8 µM and were monitored every 5 min for 3 h (37 timepoints) at a gain setting of 100. The linear region used for v_0_was the first three timepoints (0–4 min) on glucose and a fixed 60–150 min window on xylose and mannose, applied uniformly across all six parents and all eight substrate concentrations *Variant enzymes*. Each variant was assayed against all three substrates over a uniform eight-point two-fold dilution series from 1000 µM down to 7.8 µM. Enzyme concentrations were 10 nM on glucose and 100 nM on xylose and mannose, with two exceptions: variants M13 and M21 were assayed at 100 nM on glucose because its glucose activity was too low to give a detectable signal at 10 nM. Glucose progress was monitored every 5 min for 1 h (13 timepoints) and xylose/mannose progress every 5 min for 3 h (37 timepoints) at the extended dynamic range gain setting. The linear region used to compute v_0_was tuned per enzyme–substrate pair rather than applied uniformly, because both the time to saturate the substrate and the lag before reaching steady-state varied substantially across the variant panel; the full table of (enzyme, substrate) → time window is provided in the supplementary materials. All other downstream processing — background subtraction, standard-curve conversion, triplicate averaging, substrate-inhibition fit, and slope-derived *k*_cat_/*K*_m_ — was identical to the parent pipeline.

### Fragment importance analysis

To assess which parental fragments drive substrate preference, 1,000 chimeras were sampled uniformly at random from the 6^8^ = 1,679,616 combinations defined by the six parental sequences and eight recombination fragments (duplicates and any sequence already present in the assayed-sequence database excluded). Each chimera was scored with the final-round ProteinNPT checkpoint, using the complete set of 132 assayed sequences as its labelled in-context set, yielding predicted log_1_p activities on glucose, xylose, and mannose (a1, a2, a3). Non-glucose preference was defined per chimera as S = (a2 + a3)/2 − a1, i.e. the log ratio of geometric-mean non-glucose to glucose activity. The marginal effect of parent p at fragment position i was computed as the mean S over all chimeras carrying that parental fragment, minus the grand mean of S.

### Per-residue attribution

To identify which sequence differences within each fragment contributed to the fragment-level effects shown in Fig. 4g, we decomposed each parental fragment effect into residue-level contributions. The six parental sequences were aligned separately within each of the eight recombination fragments using FAMSA with a UPGMA guide tree. For each fragment, the six aligned parental sequences were one-hot encoded by alignment position and residue identity, treating gaps as a 21st character. We then fit an independent linear regression using the six parental fragment sequences as inputs and their corresponding model-derived fragment effects as targets. Because the number of sequence features greatly exceeded the six observations, the regression is underdetermined; we therefore took the minimum-norm solution, which decomposes each fragment effect among the sequence features that distinguish the parental fragments. The eight regressions yielded 938 coefficients, one for each position–residue combination observed across all fragments. These coefficients therefore provide a residue-level attribution of the fragment effects rather than independently estimated residue effects. Because the structure is that of parent 1, each alignment column contributes the coefficient corresponding to the residue carried by parent 1 at that column. Columns at which parent 1 carries an alignment gap were skipped, leaving 479 values, one per residue of the mature parent 1 sequence. The resulting residue-level values were written to the B-factor column of an AlphaFold3 model of parent 1 (mean pLDDT 96.7) for visualization in Fig. 4h. The complete residue-level attribution values are provided in the Supplementary Materials.

## Declarations

### Funding

This research was supported by National Institutes of Health award 5R01GM150929 (to P.A.R.).

### Conflict of interest

The authors declare no competing interests.

### Code availability

All software and data analysis code is available at https://github.com/RomeroLab/PRAXIS

## Notes

https://github.com/RomeroLab/PRAXIS

### Competing Interest Statement

The authors have declared no competing interest.

