## Supplementary Information for "Learning protein function through autonomous experimental interaction"

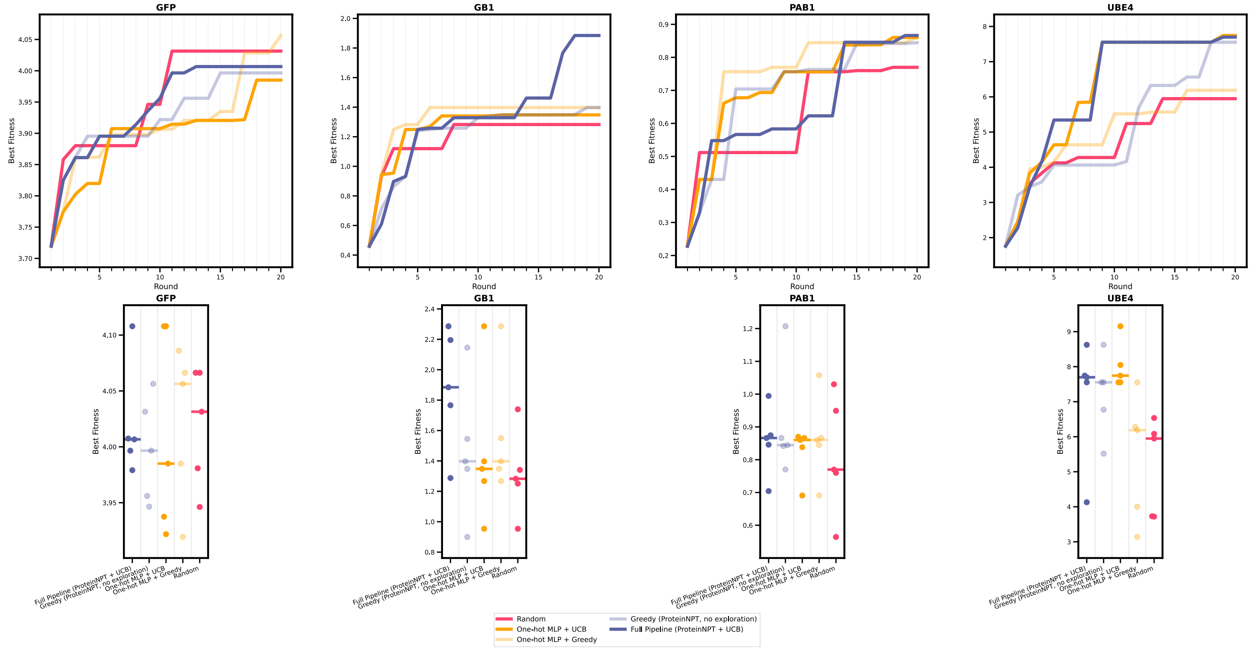


**Fig. S1:** In-silico search trajectories as shown in the main text (top row) and individual per-seed final fitness scores with a horizontal line indicating the median (bottom row). Each protein DMS dataset was searched using 5 methods x 5 random seeds.

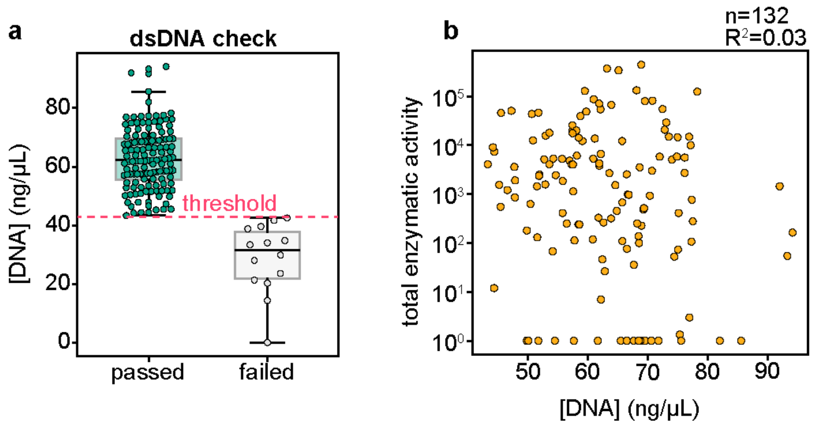


**Fig. S2:** Automated lab error handling and DNA production. (a) DNA concentration of all post-PCR gene assemblies, where the threshold determined whether sequences passed onto characterization or were added to the retry pool. (b) The relationship between the total enzymatic activity of variants and their corresponding post-PCR DNA concentration

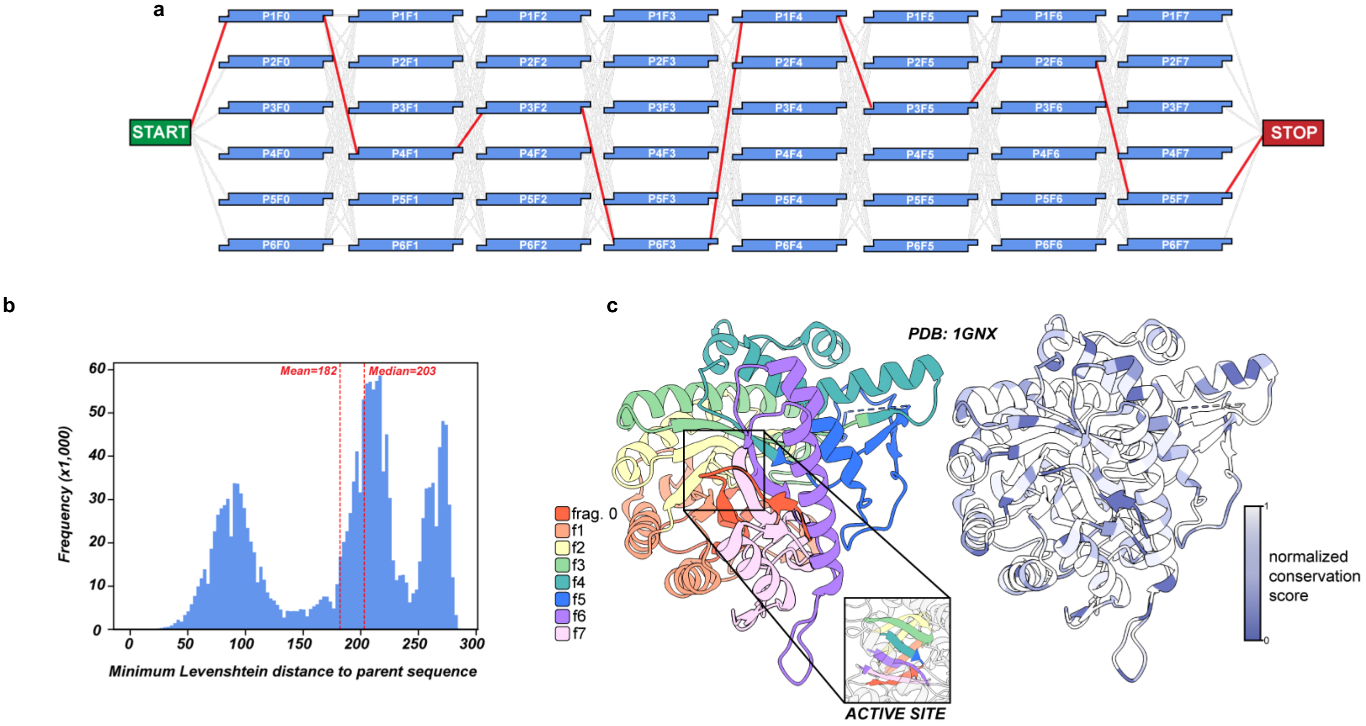


**Fig. S3:** (a) All valid enzyme chimeras can be illustrated as a path through the fragment library. There are $6^8$ = 1,679,616 possible paths from START to STOP. (b) Distribution of minimum Levenshtein (edit) distances of all ~1.7 million possible enzyme chimeras to the nearest parent enzyme. (c) Locations of the eight fragments (left) and amino acid conservation overlaid on the GH1 structure (right). Fragments encompass distinct regions of the enzyme structure and each contribute a beta strand to the inner beta barrel. Chimeric assembly results in a diverse set of unique variants.

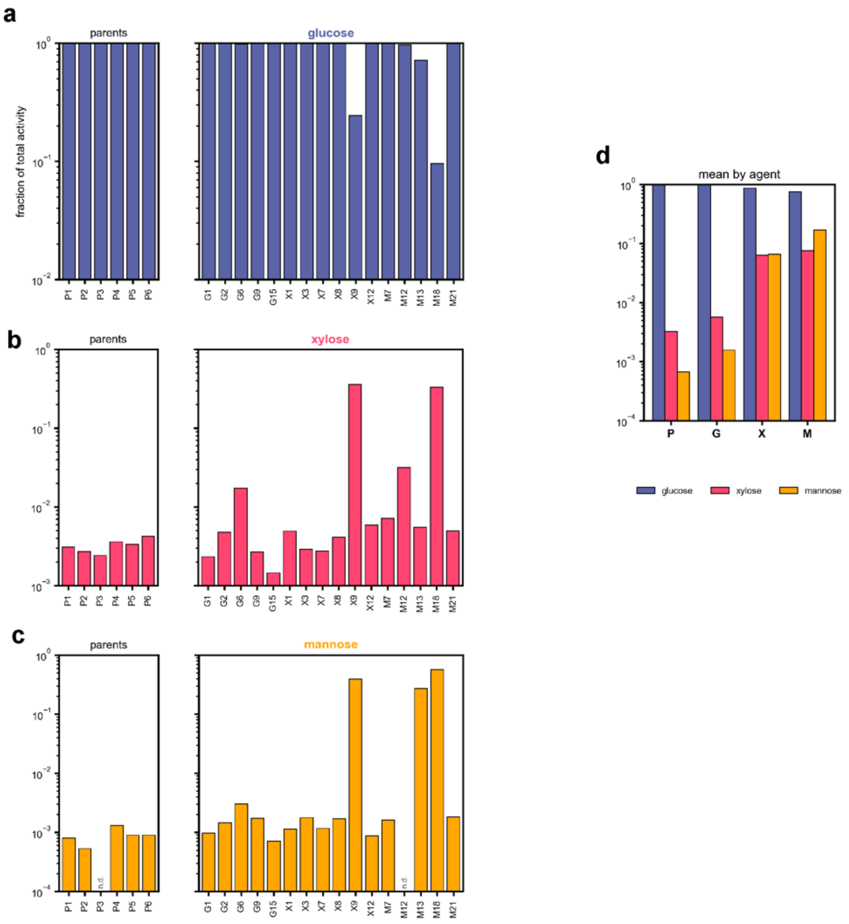


**Fig. S4:** A selected set of variants from the autonomous engineering campaign were selected for manual testing, where the autonomous experimental conditions were replicated and conducted by a human scientist. Each variant (x-axis) was tested against all three substrates, the initial slope of the RFU/time curve was converted to [S]/time by interpolation with a standard curve of 4-methylumbelliferone at known concentrations. Grey shading indicates the region extracted for slope calculation. The fraction of total activity (y-axis) was calculated from the activities on each of the three substrates, e.g. $a_1 / (a_1 + a_2 + a_3)$.

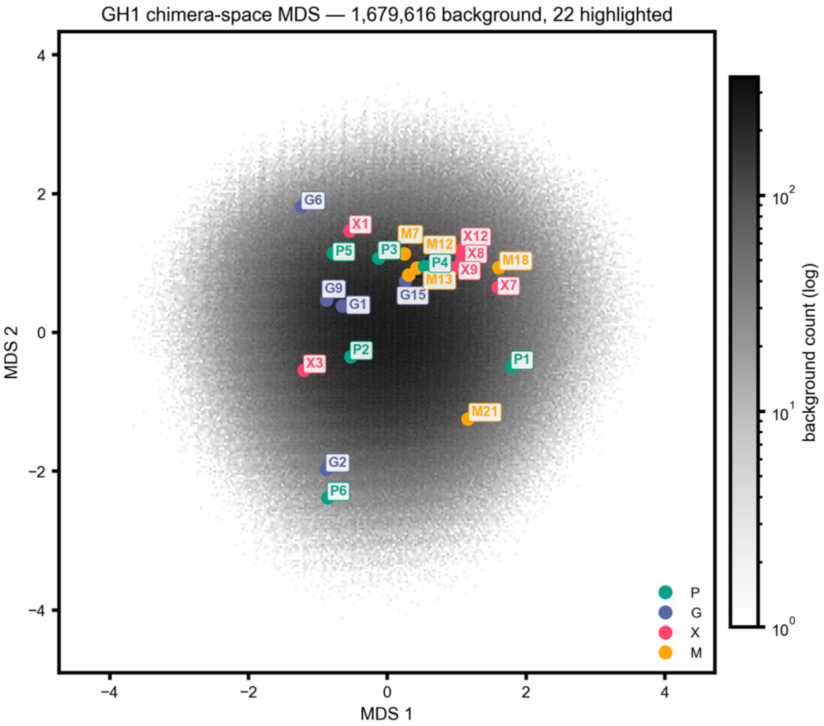


**Fig. S5:** Multidimensional scaling (MDS) of enzyme chimeras. The entire chimeric assembly space is plotted as greyscale, and the parents and tested variants are show in colors. The enzymes discovered by the non-native agents (agents X and M) occupy similar regions of chimera space.

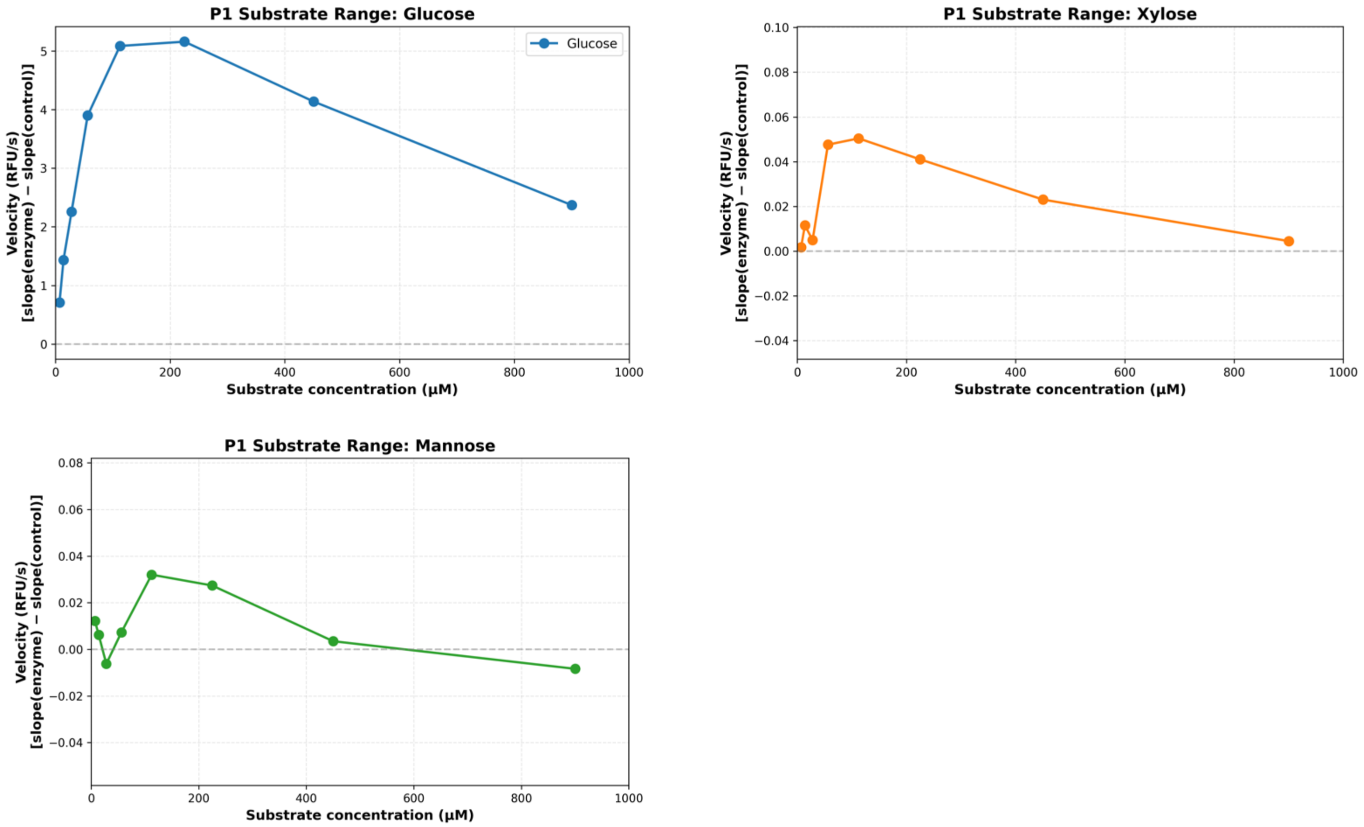


**Fig. S6:** Pilot lysate assays of Parent 1 against all three sugar substrates showing clear inhibition of enzyme activity at high [S], used to determine optimal substrate concentration for automated experimentation.

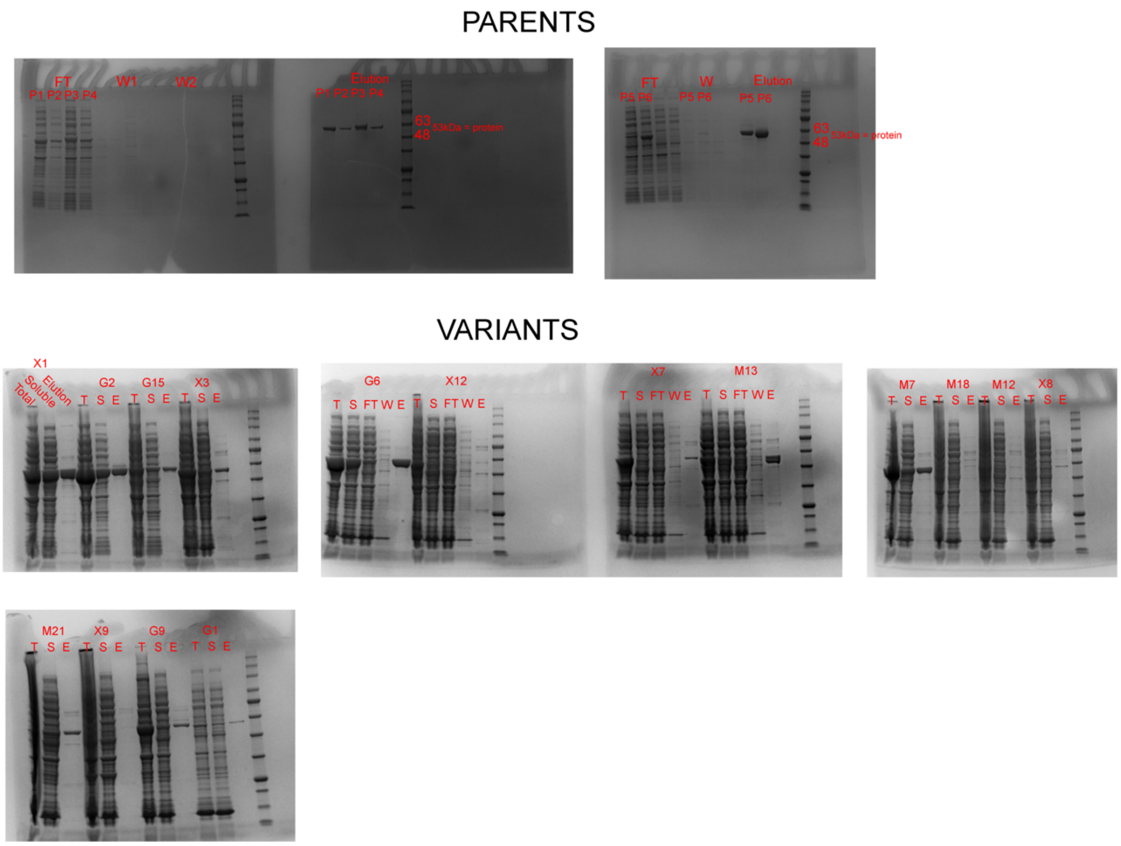


**Fig. S7:** SDS-PAGE images of the purifications and corresponding eluted fractions of all parent enzymes and tested variants.

**Table S1:** ProteinNPT hyperparameters used in PRAXIS model training and sequence acquisition.

| **ProteinNPT Hyperparameters** | | |
|---|---|---|
| Component | Hyperparameter | Value |
| **Architecture** | Protein encoder | ProteinNPT (5 layers) |
| | Embedding source | ESM2-650M (frozen) |
| | Internal embedding dim | 200 |
| | Self-attention heads | 4 |
| | FFN hidden dim | 400 |
| **Training** | Total training steps (per BO round) | 5,000 |
| | Batch size (assay sequences / GPU) | 425 |
| | Optimizer | AdamW |
| | LR schedule | Warmup (100 steps) $\rightarrow$ cosine decay |
| | Max / min learning rate | $3 \times 10^{-4}$ / $1 \times 10^{-5}$ |
| | Adam $(\beta_1, \beta_2)$ / $\epsilon$ | (0.9, 0.999) / $10^{-8}$ |
| | Gradient accumulation | 1 |
| | Gradient clipping | 1.0 |
| | Weight decay | $5 \times 10^{-3}$ |
| | FP16 training | Enabled |
| | Indel mode | Enabled |
| **Regularization** | Attention dropout | 0.1 |
| | Activation dropout | 0.1 |
| | Token dropout | 0.0 |
| | Label smoothing | 0.0 |
| | Seed | 2023 |
| **Targets & Loss** | Primary objective | Specificity-weighted activity (agent-specific) |
| | Auxiliary labels | Tranception zero-shot score; binary activity flag |
| | Loss weighting | Annealed AA reconstruction + target MSE |
| **Uncertainty & Acquisition** | Uncertainty estimation | MC dropout (5 samples) |
| | Acquisition rule | UCB: $\mu(x) + \alpha\sigma(x)$, $\alpha = 1$ |
| | Diversity enforcement | ESM2-650M embeddings + $k$-means clustering |
| **Sampling Loop** | Sampling mode | Conditional fragment resampling |
| | Parents per round | Top quartile by specificity objective |
| | Offspring per inner batch | 30 conditional sequences |
| | Total samples per iteration | 1,000 |
| | BO rounds | 25 |
| | Acquisition batch size | 2 sequences / agent / round |
